# Myoglobin leaching into the serum of IDA mice is driven by the high activation of sGC under anemic conditions which induces myoglobin expression

**DOI:** 10.1101/2025.10.03.680120

**Authors:** Arnab Ghosh, Nupur K. Das, Mamta P. Sumi, Blair Tupta, Chaitali Ghosh, Dennis J. Stuehr, Yatrik M. Shah

**Affiliations:** Department of Inflammation and Immunity, Cleveland Clinic Research, The Cleveland Clinic, Cleveland, Ohio-44196; Department of Biomedical Engineering, Cleveland Clinic Research, The Cleveland Clinic, Cleveland, Ohio-44196, USA; Department of Molecular and Integrative Physiology, University of Michigan, Ann Arbor, Michigan 48109

**Keywords:** Anemia, Erthropoiesis, Heme-dependent, Heme-maturation, Myoglobin

## Abstract

Our study reveals that status of the sGC heterodimer or its subsequent activation aligns with active erythropoiesis, and this heterodimer also correlates with the expression of myoglobin (Mb) or HO1. In this study we found that Mb expression which is driven by iron restriction and high sGC activation in iron deficiency anemia (IDA) leaches out more into the serum relative to non-anemic WTs. Tissues from IDA mice of both models developed either by nutritional iron deprivation or by ablation of ferroportin (Fpn) gene or from iron refractory iron deficiency anemia (IRIDA) mice found that Mb expression follows a variable pattern in different tissues but always correlates to the status of the sGC heterodimer or its subsequent activation. Here higher Mb expression happening in anemic (IDA, Fe<5 ppm or IDA, Fpn) or non-anemic WT mice is both due to iron restriction and an elevated sGC heterodimer that corroborated with greater sGC activation. More importantly we find significant leaching of Mb into the serum of these anemic (IDA) mice from both models and our spectral data suggests that this Mb is heme-free. This Mb leaching in anemia is a cumulative impact of Mb secreting out from various tissues including lungs, spleen, skeletal or cardiac muscles where Mb is expressed and not just in the skeletal muscles where Mb expression is low. Based on these findings we construct a working model of anemia, where high activation of sGC under anemic conditions (Fpn ablation or restricted Fe diet) induces Apo-Mb or heme-free Mb expression which can then leach out into the serum. Our findings of Mb leaching are novel and can find further application as a diagnostic strategy in anemia.

## INTRODUCTION

Nitric oxide (NO) is a signal molecule and plays a critical role in the regulation of vascular tone (1-4), displays anti-platelet (4-8) and anti-inflammatory properties (9). NO generated by NOS enzymes (10-13) or produced by noncanonical means (14), is a central component of the NO-sGC-cGMP signal pathway and works by activating the soluble guanylate cyclase (sGC), an obligate heterodimer of α-subunit (sGCα) and β-subunits (sGCβ) (15-18). Our studies found that low doses of NO can directly trigger heme-insertion in muscle Mb or trigger Hbαβ heterodimerization (19, 20) and as these events imply that NO-sGC signaling is involved in globin maturation we need to determine how this signal pathway can be used to predict the state of erythropoiesis (21) or muscle Mb build up.

Iron deficiency anemia occurs when the level of healthy RBCs decreases and is the most common disease affecting approximately 5 million people in the United States (22). Previous findings suggest that iron restriction increases Mb expression (23), while our study indicates that sGC plays a critical role in Mb heme-maturation (24), which under anemic conditions may suggest an increase in heme-free Mb or Apo-Mb. While there is evidence of Mb breakdown from the muscles under anemia (25), we speculated that an increase in apo-Mb which otherwise has no cellular function may result in Mb leaching out in the blood. In the current study we used expression analysis and IP assays in erythropoietic tissue supernatants to first predict the status of the labile sGC heterodimer (18) and determine whether its NO heme-dependent activation aligns with erythropoiesis (21). We then used tissue supernatants from two different anemic (IDA) mice models (one generated by low Fe fed diet and the other by Fpn deletion) and one from IRIDA (made by deletion of transmembrane serine protease 6) (26, 27) mice to determine whether the status of the sGC heterodimer correlates with its corresponding tissue Mb expression. Serum analysis was then done to determine the extent of Mb leaching in anemic mice relative to non-anemic WTs. Our findings suggest that status of the labile sGC heterodimer aligns with erythropoiesis and to the expression of Mb in anemia, while significant amounts of Mb leaches out in the serum during anemia.

## MATERIAL AND METHODS

### Reagents

All chemicals were purchased from Sigma (St. Louis, MO) and Fischer chemicals (New Jersey). NO donor, SNAP (S-nitroso-N-acetyl-D,L-penicillamine), Phosphodiesterase inhibitor 3-isobutyl-1-methylxanthine (IBMX) sGC inhibitor ODQ ([1H-[1,2,4]oxadiazolo-[4, 3-a]quinoxalin-1-one]) and sGC stimulator BAY 41-2272 (BAY-41, heme-dependent) were purchased from Sigma. Bone marrow (BM) and spleen of 3, 5 or 10 week old mice (C57BL/6) or neonatal liver from 2-9 day mice were purchased from Jackson Labs (Bar Harbor, ME, USA), while liver, spleen, skeletal muscles, hearts, lungs or serum samples from TMPRSS6 or IDA (Fe < 5ppm and Fe 350 ppm) and control WT mice were obtained from Dr. Yatrik Shah’s lab (Univ. of Michigan). cGMP ELISA assay kits were obtained from Cell Signaling Technology (Danvers, MA, USA). Smooth muscle cells and endothelial cells specific culture media was purchased from Lonza (Basel, Switzerland). Protein G-sepharose beads were purchased from Sigma and molecular mass markers were purchased from Bio-Rad (Hercules, CA, USA).

### Antibodies

Antibodies were purchased from different sources. Supplemental Table S1 describes various types of antibodies used and its source.

### Culture and differentiation of C2C12 cells

For C2C12 culture, the myoblasts were induced to differentiate into myotubes between 0 and 96 h by growing in media containing 2% horse serum, with media changed every 48 h (24). To investigate the effects of sGC inhibition on myoblast differentiation and Mb heme-isertion, sGC inhibitor ODQ (1µM) (28) was added to the differentiation media between 24-96 h. The cells were imaged every 24 h before being harvested. In all cases wherever applicable cell supernatants were assayed for protein expression by western blot, binding assays by IPs and depiction of Mb heme-maturation by heme staining (24, 29), as indicated.

### Anemic Mice generation

Iron deficiency anemia (IDA) was induced in mice either genetically or by nutritional iron (Fe) deprivation, while iron refractory iron deficiency anemia (IRIDA) mice was developed genetically. For inducing anemia by nutritional iron (Fe) deprivation 4-6 weeks old C57Bl6/J mice were fed with AIN93G rodent diet supplemented with <5 ppm (iron-restricted diet) relative to control 350 ppm iron enriched diet for 4 weeks following procedures as previously described (30). The other IDA mice was developed genetically by ablation of the intestinal ferroportin (Fpn) gene (Intestinal Fpn-KO). In this a tamoxifen inducible, intestinal epithelium specific Cre recombinase (*Vil*^CreERT2^) were used in combination with homozygous floxed mice for ferroportin (*Fpn*^fl/fl^) to generate intestinal Fpn1 KO (*Fpn1*^ΔIE^: *Vil*^CreERT2^; *Fpn*^fl/fl^) mice. Following tamoxifen injection (100 mg/kg IP, 5 days), mice were maintained on regular rodent diet containing 300 ppm iron for 12 weeks. Cre negative *Fpn*^fl/fl^ mice served as controls (31, 32). TMPRSS6 KO mice model for IRIDA was generated as described earlier (26, 27).

### Western blots and Immunoprecipitations (IPs)

Cell or tissue supernatants generated from mice lung, skeletal muscles, liver, spleen, hearts or BM were analyzed for protein expression by western blots or for protein-protein interactions by immunoprecipitation assays (IPs) (33). For western blots 50-80ug of the tissue supernatants were run on SDS-PAGE (8 or 15%), transferred to the same PVDF membrane and probed with a specific antibody. β-actin was used as a loading control. Multiple protein detection was achieved by stripping the membranes and re-probing with specific antibodies. For immunoprecipitations (IPs), 400-600 μg of the total tissue supernatant was precleared with 20 µl of protein G-sepharose beads (Amersham) for 1 h at 4º C, beads were pelleted, and the supernatants incubated overnight at 4 ºC with 3 μg of the indicated antibody. Protein G-sepharose beads (20 µ L) were then added and incubated for 1 h at 4 ºC. The beads were micro-centrifuged (6000 rpm), washed three times with wash buffer (50 mM HEPES pH 7.6, 100 mM NaCl, 1 m(34)M EDTA and 0.5% NP-40) and then boiled with SDS-buffer and centrifuged. The supernatants were then loaded on SDS-PAGE gels and western blotted with specific antibodies. Band intensities on westerns were quantified using Image J quantification software (NIH).

### Nitrite in the culture media

Nitrite was measured using ozone-based chemiluminescence with the triiodide method and using the Sievers NO analyzer (GE Analytical Instruments, Boulder, CO, USA) (34, 35).

### cGMP enzyme-linked immunosorbent assay

sGC enzymatic activity in the mice tissue supernatants was determined by adding 250 μM GTP and 50 µM SNAP plus 10 μM of sGC activators BAY 41-2272 and incubating for 10 min at 37 °C. Reactions were quenched by addition of 10 mM Na_2_CO_3_ and Zn (CH_3_CO_3_)_2_. The tissue supernatants were passed through the desalting PD-spin trap G-25 columns (GE Healthcare) prior to addition of the above constituents. The cGMP concentrations was then determined by ELISA and was a measure of sGC activity in the tissue supernatants (18).

### UV-Visible spectral measures

UV-visible absorption spectra of mice serum samples were recorded at room temperature between 350-700 nm on a Shimadzu spectrophotometer. Equal amounts of total protein supernatants were used for respective wavelength scans. The heme content for Hb was determined from the Soret absorption peak at 414 nm, using the extinction coefficient of 342500 M^-1^cm^-1^ (414 nm) and a manipulation to account for the variable absorbance contributions that were attributable to sample turbidity. This involved creating a baseline for each scan by drawing a line that connected the absorbance values at 380 and 470 nm. The additional Soret absorbance at 414 nm above this baseline was then used to calculate the heme content of Hb (36).

## RESULTS

### Status of NOS enzymes, HO-1 and sGC heterodimer during erythropoiesis

To assess the involvement of the NO-sGC signal pathway during erythropoiesis we determined the expression of NOSs and sGC in the bone marrow (BM), spleen or liver tissues, as these are the three vital erythropoietic locations in mammals (21). Heme oxygenase 1 (HO1) expression was also determined as it is known to be upregulated during erythropoiesis (37-39). Doing western blots in the tissue supernatants of 3, 5, 10 week or 2, 3, 5 and 9 day old mice (Figs. 1A, D), we found predominant stable expression of iNOS and sGCβ1, while the HO1 expression increased in the BM and declined in the spleen. IP assays showed the labile sGC heterodimer to increase in the BM of 10 week old mice where the erythropoietic activity is known to surge and is sufficient to sustain steady-state adult erythropoiesis (21) (Fig. 1B). The status of the sGC heterodimer was found to decline in the spleen or liver where fetal erythropoiesis is lost within 1 or 7 weeks respectively (21) (Fig. 1B, E). Also expression of HO1 paralleled the status of the sGC heterodimer in BM and spleen (Fig. 1B), which suggests that the HO1 expression and status of the sGC heterodimer are synchronous with the rise and fall of the reported erythropoietic activities (21). This was further corroborated by EPO expression in BM or spleen (Fig. 1B) which was also aligned to the erythropoietic activities (21). Both the sGC activation with NO and BAY-41 and the basal sGC activity (Fig. 1C) also correlated with the status of the sGC heterodimer. Together these data suggests that HO1 expression aligns with the status of the sGC heterodimer and these can be used as molecular signatures to track erythropoiesis.

**Figure 1.**
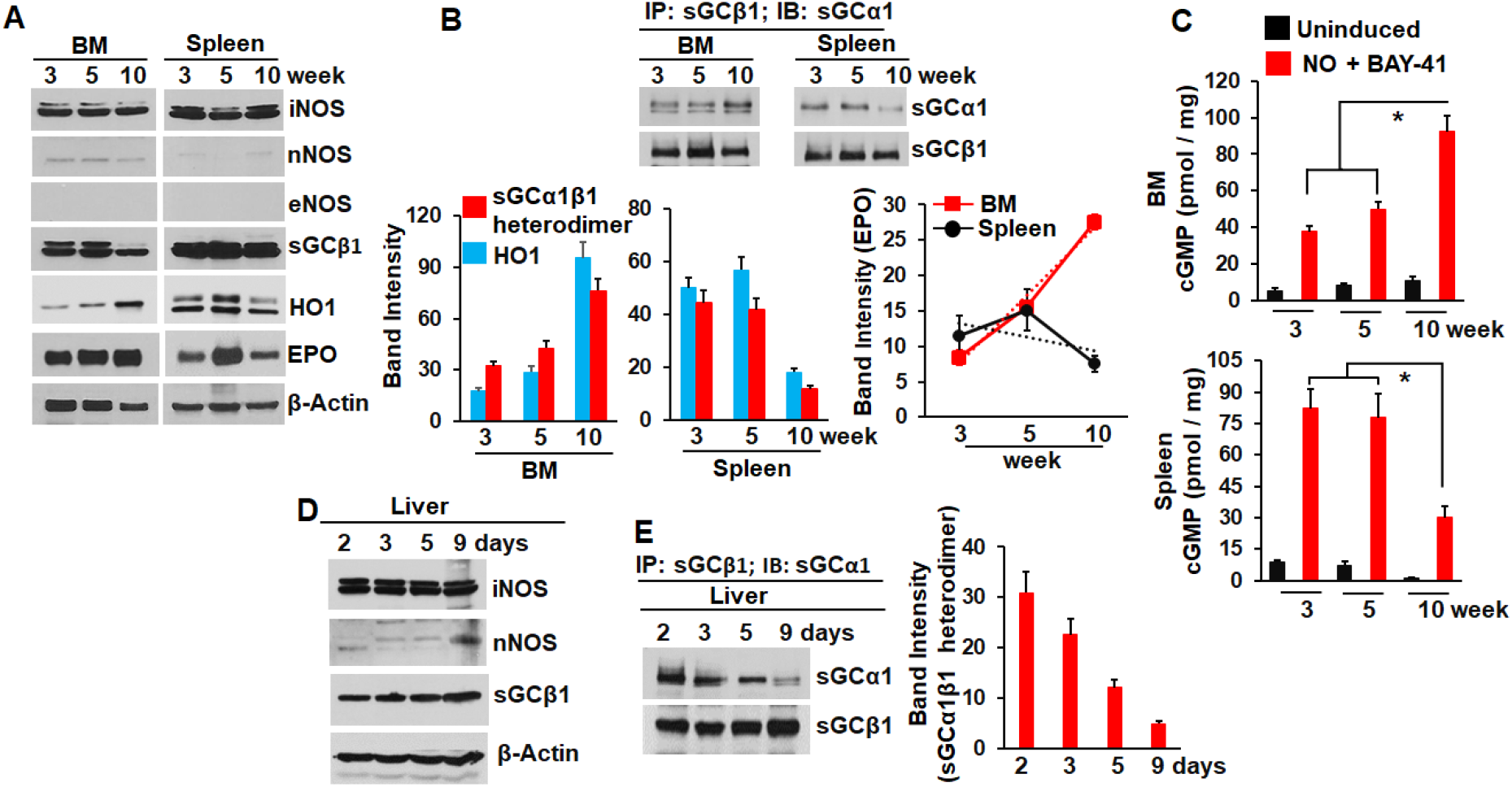
Status of sGC heterodimer, HO1 and NOSs during erythropoiesis. Tissue supernatants of days/ weeks old mice was prepared from its corresponding bone marrow (BM), spleen or liver and used in western blots, IPs or sGC activation assays. (A, B) Expression of indicated proteins with β-actin used as a loading control and status of the sGC heterodimer (IP) or HO1 & EPO expression (band intensities from A or B) in the BM and spleen supernatants of 3, 5 & 10 week old mice. (C) cGMP estimation by ELISA, determined from sGC activation with NO and/ BAY-41 or the basal sGC activity. (D, E) Expression of indicated proteins and status of the sGC heterodimer (IP & its band intensity) in the liver supernatants of 2, 3, 5 and 9 day old mice. All depicted values are mean of n=3 repeats ± SD. *p < 0.05, by an independent student’s *t*-test.

### Status of sGC heterodimer aligns with tissue Mb expression and is sensitive to changes in Fe levels in anemic mice

To determine how anemia may impact the status of the sGC heterodimer, we used tissues from TMRSS6 mice causing iron refractory iron deficiency anemia (IRIDA) (26) or from iron deficiency anemic (IDA) mice developed by feeding a low iron diet (Fe <5ppm, nutritional deprivation of iron) relative to non-anemic control mice kept on an iron-replete diet (Fe 350ppm) (30). Western blots on tissue supernatants showed presence of iNOS, sGCβ1 along with varying HO1 levels that correlated with tissue Myoglobin (Mb) levels (Fig. 2A, B). IPs done on these tissue supernatants showed the sGC heterodimer to be more in the TMPRSS6 mice relative to WT in the liver and comparable in the spleen (Fig. 2B). Interestingly the sGC heterodimer status fluctuated with the Fe levels of the IDA or control non-anemic mice tissues and displayed contrasting states in the liver vs spleen (Fig. 2B). Both the Mb and HO1 levels were significant in the spleen but lowered in the liver (Fig. 2A) and were synchronous with the changes in sGC heterodimer which was sensitive to changes in the Fe levels (<5 vs 350ppm, Fig. 2B). Since sGC is needed for heme-maturation of Mb (24), a direct correlation of the sGC heterodimer status with tissue Mb expression authenticates our previous findings. Doing sGC activation assays on spleen supernatants with NO and BAY-41 corroborated the status of the sGC heterodimer (Fig. 2C). Together these data imply that the sGC heterodimer, is sensitive to changes in Fe levels and can be tracked by following expression of Mb or HO1.

**Figure 2.**
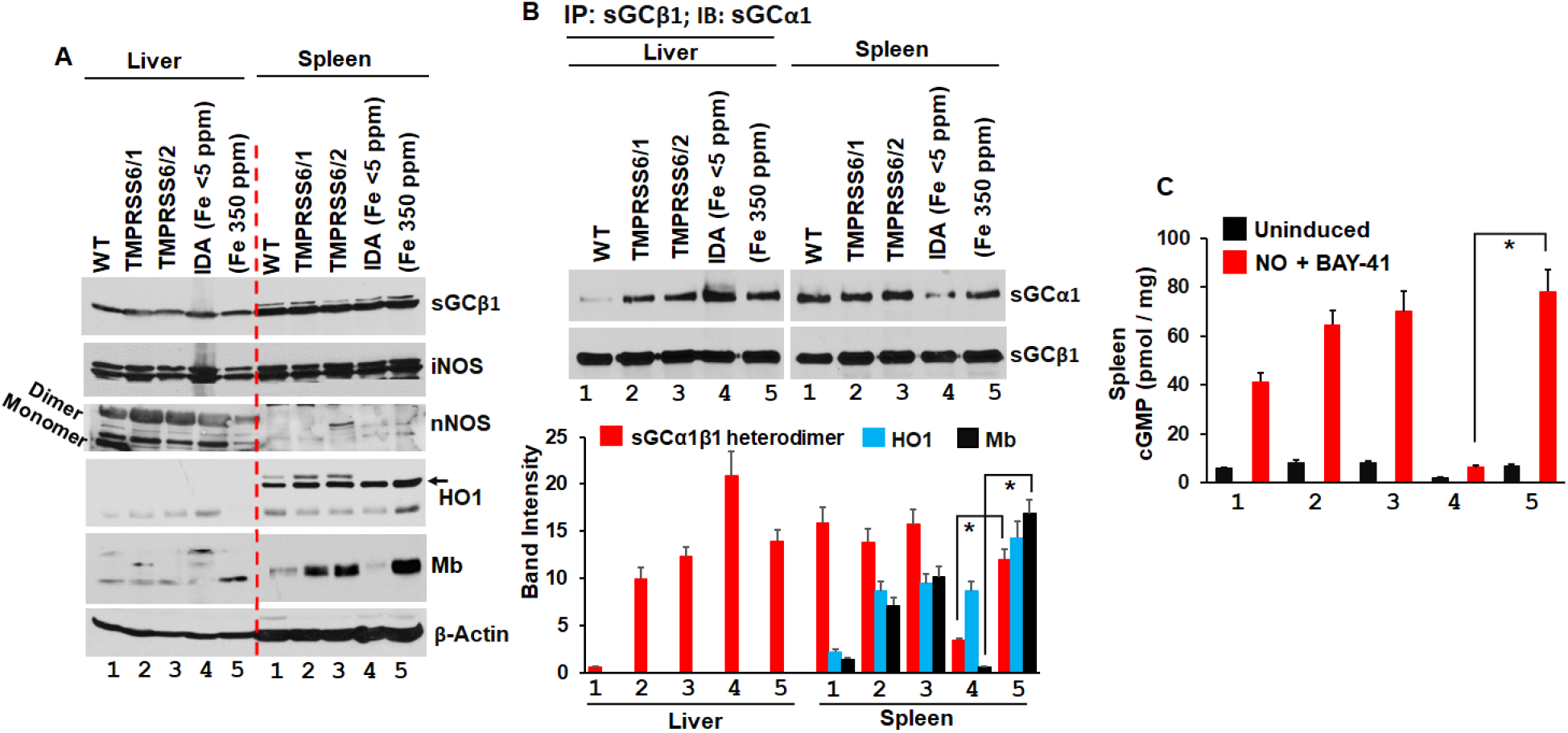
Status of sGC heterodimer correlates with HO1 and tissue Mb expression in the spleen. Liver and spleen supernatants prepared from WTs or anemic mice were used for western blots or sGC activity assays. (A) Protein expression by western blots as indicated with β-actin used as a loading control. (B) IPs depicting the status of sGC heterodimer in TMPRSS6 (groups 1/2) or IDA with two different Fe diet as indicated. Band intensities (mean of n=3 mice tissues) depicting relative correlations with sGC heterodimer status, HO1 and Mb expressions between WT or anemic mice tissues as indicated. (C) cGMP estimation by ELISA, determined from sGC activation with NO and/ BAY-41 as indicated. All depicted values are mean of n=3 repeats ± SD. *p < 0.05, by student’s unpaired *t*-test.

### Mb from anemic IDA mice leaches out more into the serum relative to WTs

As in IDA mice developed by restricted Fe (<5ppm) diet, Mb expression can be induced (23) which is predominantly heme-free Mb or apo-Mb and this coupled with the higher activation of sGC (24) as found in the spleen supernatants can form greater holo-Mb protein when supplemented with Fe (350ppm) (Figs. 2A-C). Testing Mb expression in the liver/spleen tissues we found a variable pattern of Mb expression (Fig. 3A) and it seemed to be regulated by the status of the sGC heterodimer as Fig. 3A correlates to Figs. 2B, C. Here higher Mb expression in the liver/spleen happening in anemic (IDA, Fe<5 ppm) or non-anemic (Fe 350 ppm) mice seems to be due to an elevated sGC heterodimer which indicates greater sGC activation (Fig. 2C). As Mb cannot be a product of RBC lysis, we probed for Mb in the serum and found greater amounts of Mb protein in the serum samples of IDA (Fe <5ppm) mice relative to non-anemic (Fe 350ppm) mice (Fig. 3B). This suggested a higher Mb leaching out into the serum of IDA mice, relative to non-anemic mice. While spectral data of serum was masked by hemoglobin soret (414 nm) (Fig. 3B), it also indicated that leached out Mb maybe heme-free which does not have a function and so is secreted out into the blood. Together our data indicates that a high Mb expression in the tissues directly correlates with a more active sGC and that more Mb leaches out in the serum of IDA mice relative to non-anemic mice.

**Figure 3.**
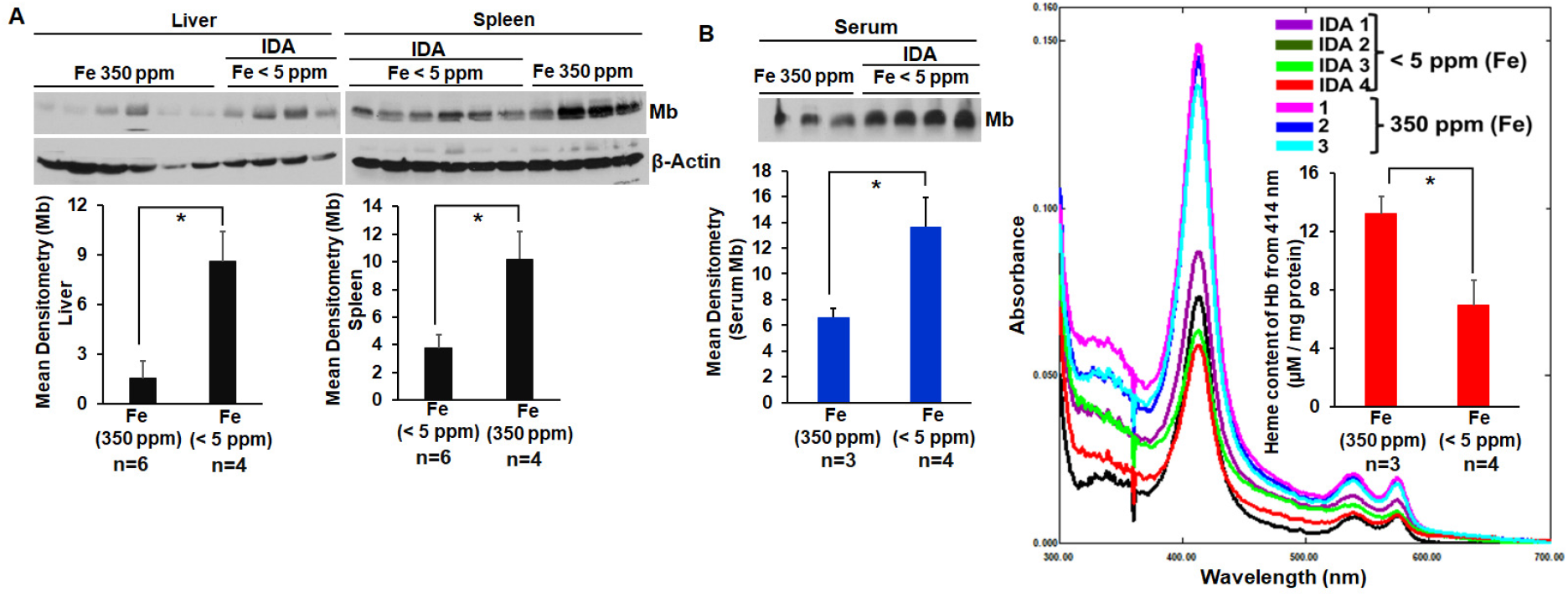
Myoglobin from anemic (IDA) mice developed by feeding low Fe diet (<5 ppm) leaches out more into the serum relative to high Fe diet mice. Liver and spleen supernatants prepared from anemic mice (IDA) fed low Fe (<5 ppm) or non-anemic mice fed high Fe (350ppm) diet and corresponding serum samples were used for western blots or UV-visible spectra. (A) Western blots depicting Mb expression with β-actin as a loading control in tissue or serum samples with corresponding Mb densitometries as indicated. (B) Heme-spectra of the serum samples from anemic mice (IDA) fed low Fe (<5 ppm) or non-anemic mice fed high Fe (350ppm) and calculated heme-content of Hb as indicated. All depicted values are mean of n=3 or 4 for IDA (Fe <5 ppm) or non-anemic (Fe 350ppm) mice samples as indicated ± SD. *p < 0.05, by student’s unpaired *t*-test.

### Contrasting patterns of Mb expression in skeletal and cardiac muscles in anemic mice correlates to the status of sGC heterodimer, while Mb leaching in the serum also occurs in a genetic model of anemia

We then used a genetic model of anemia (IDA, Fpn) whereby ablation of Ferroportin (Fpn) gene in the intestinal epithelium induces anemia in mice. Using skeletal and cardiac muscle tissues from IDA (Fpn) mice, we first compared Mb expression relative to WTs. As depicted in Figs. 4A, B there was contrasting pattern of Mb expression in the skeletal muscles (depleted in Mb) relative to cardiac tissues (elevated in Mb) in the anemic mice and these in each case directly correlated to the status of the sGC heterodimer (Figs. 4A, B & D, E), which directly corroborated to the activation of sGC (Fig. 4F). Additionally the sGC heterodimer status correlated directly with not only the Mb expression but also with the Mb heme as we determined from differentiation of C2C12 skeletal muscle myoblasts (Fig. S1), where the NO-sGC pathway is involved and inhibiting sGC with its inhibitor ODQ (28), blocked both the sGC heterodimerization and Mb heme-insertion, with the subsequent stalling of differentiation (Fig. S1) (24). These results imply that both apo-Mb (heme-free) induction and holo-Mb (heme-containing) formation may depend on an active sGC. We also found increased leaching or secretion of Mb into the serum of these anemic mice (IDA, Fpn) relative to WTs (Fig. 4C), which suggests that Mb leaching maybe a common phenomenon in anemia as it occurred in both IDA models, either caused by nutritional Fe deprivation (Fe<5 ppm) or by genetic ablation of Fpn gene. The serum heme spectra was predominant for the heme soret of Hb (414 nm) (Fig. 4G), while masking the Mb heme as noted earlier (Fig. 3B). However the serum heme spectra of Hb was significantly low in heme content for IDA (Fpn) mice which also suggests that leached Mb heme in anemic mice is heme-free relative to WTs. Moreover there was no shift in the heme soret towards Mb heme (409 nm) (24) even when the Hb soret was very low in IDA (Fig. 4G), which further indicates that the leached Mb is heme-free. Testing for other tissues like the lungs from the Fe deprived anemic (IDA) mice, also showed elevation in Mb levels relative to non-anemic high Fe diet mice (Fig. 5A) which directly correlated to an elevated sGC heterodimer (Fig. 5B), suggesting that it was similar to the cardiac tissues of IDA (Fpn) mice, further indicating that both nutritional and genetic models may have physiological semblance. Based on these findings we construct a working model of anemia (Fig. 6), where Fe deprivation cause by Fpn ablation or restricted Fe diet, along with an activated sGC can induce Mb expression in anemia. This Mb leaches out into the serum in anemia and is probably heme-free. This coupled by a higher sGC activation under anemic (IDA) conditions can cause the apo-Mb protein to become an active receptor for heme protoporphyrin and if Fe supplementation is provided it can form the holo-Mb protein. However more importantly our findings of Mb leaching are novel and can find further application as a diagnostic strategy in IDA.

**Figure 4.**
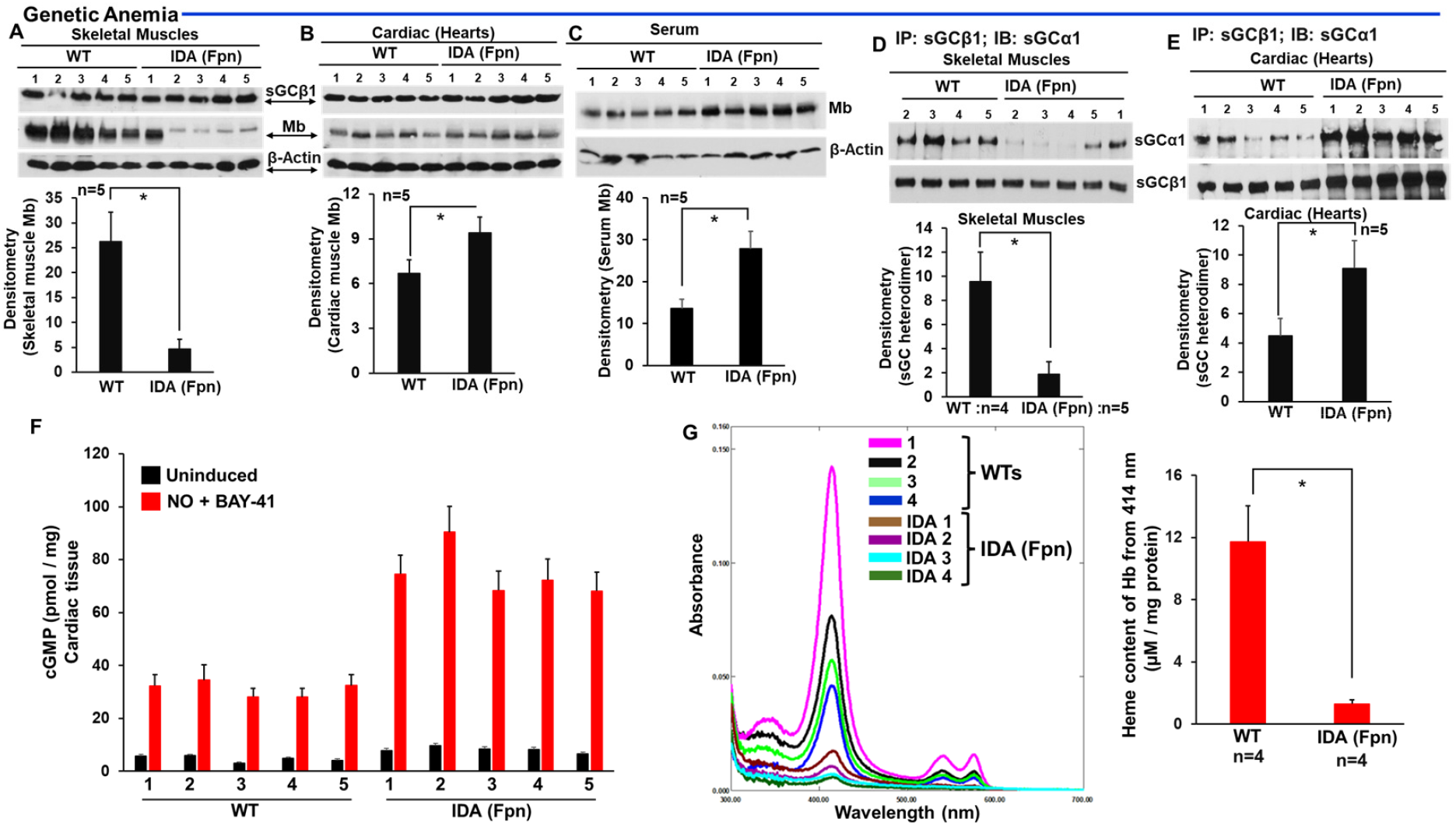
Mb leaching into the serum also occurs in a genetic model of anemia (IDA, Fpn) and contrasting patterns of Mb expression in anemic mice tissues correlates to the status of sGC heterodimer. Skeletal and cardiac muscle tissue supernatants prepared from WTs or anemic (IDA, Fpn) mice created by ablation of Ferroportin (Fpn) were used for western blots, IPs and sGC activity assays. (A, B, C) Protein expression by western blots as indicated with β-actin used as a loading control and corresponding densitometries for Mb expression. (D, E) IPs depicting the status of sGC heterodimer in IDA mice tissues relative to WTs and corresponding densitometries for sGC heterodimer. Values are mean of n=4 or 5 from IDA (Fpn) or WT samples as indicated ± SD. (F) cGMP estimation by ELISA, determined from sGC activation with NO + BAY-41 as indicated. All depicted values are mean of n=3 repeats ± SD. (G) Heme-spectra of serum samples from anemic mice (IDA, Fpn) or WTs, and calculated heme-content of Hb as indicated. All depicted values are mean of n=4 for IDA (Fpn) or non-anemic WT mice samples as indicated ± SD. *p < 0.05, by student’s unpaired *t*-test.

**Figure 5.**
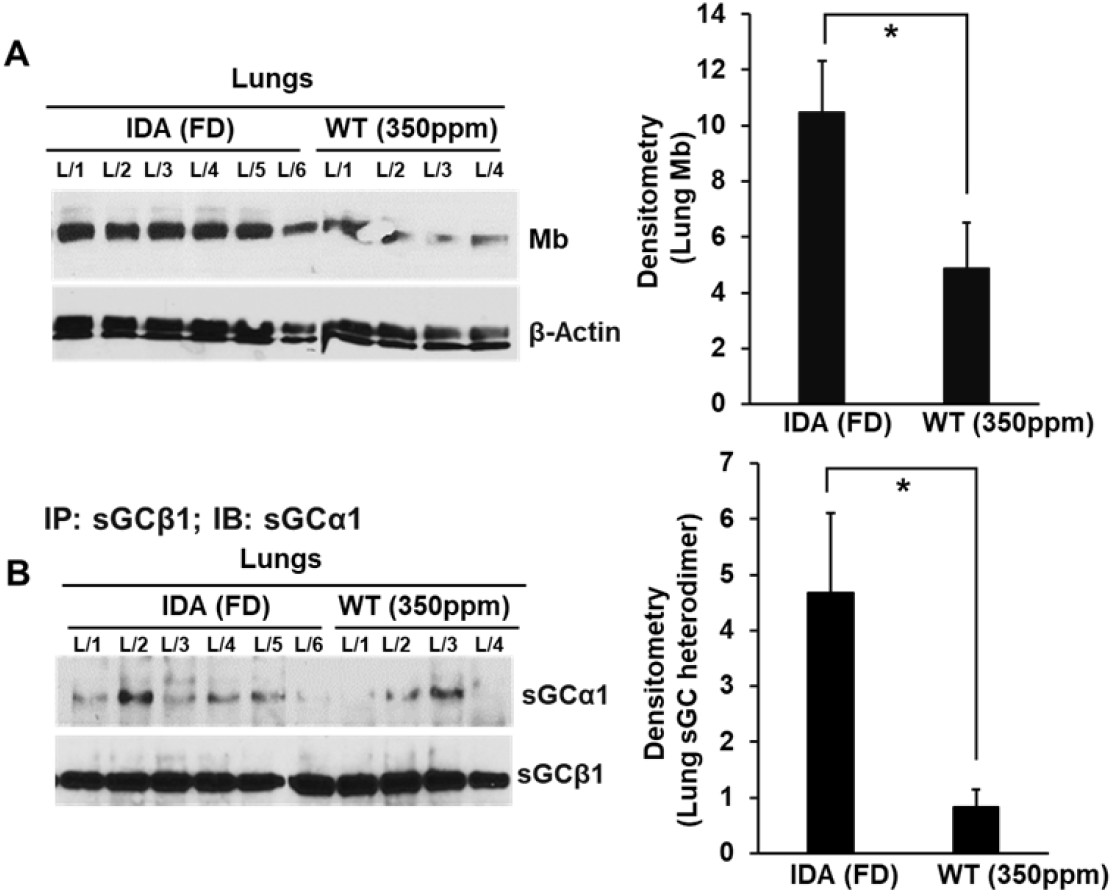
Mb expression in the lung tissues of IDA (Fe <5 ppm) mice is driven by the status of the sGC heterodimer. Lung tissue supernatants prepared from non-anemic (Fe 350 ppm) or anemic IDA (Fe <5 ppm) mice were used for western blots and IPs. (A) Protein expression by western blots as indicated with β-actin used as a loading control and corresponding densitometries for Mb expression. (B) IPs depicting the status of sGC heterodimer in IDA mice lungs relative to non-anemics and corresponding densitometries for sGC heterodimer. All depicted values are mean of n=5 or 4 for IDA (Fe <5 ppm) or non-anemic (Fe 350ppm) mice samples as indicated ± SD. *p < 0.05, by student’s unpaired *t*-test.

**Figure 6.**
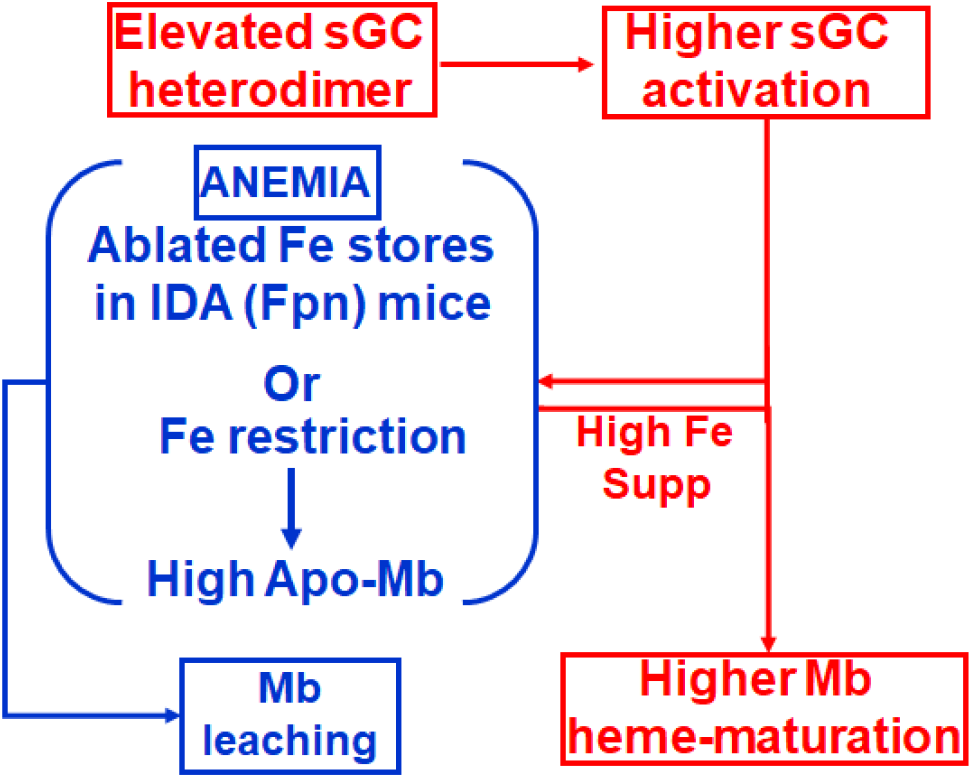
Flow diagram depicting mechanism of Mb induction in anemia and its subsequent leaching into the serum. IDA developed by Fe deprivation either caused by low Fe diet (<5 ppm) or by genetic ablation of Fpn can induce apo-Mb. This coupled by a higher sGC activation under anemic (IDA) conditions can cause the apo-Mb protein to become an active receptor for heme protoporphyrin + Fe to form holo-Mb protein. This Mb is leached/secreted out into the serum of anemic mice.

## DISCUSSION

Our study finds that the status of sGC heterodimer or its subsequent activation aligns with active erythropoiesis, and also directly correlates with the expression of Mb or HO1 in various organs where these proteins are expressed (Figs. 1 and 2). Using myoblast differentiation of skeletal muscle cells (C2C12) we further demonstrate that Mb heme-insertion is dependent on an active sGC heterodimer, abrogation of which inhibits Mb heme-maturation (Fig. S1), implying that both Mb protein expression and its post-translational heme-maturation are determined by the status of the sGC heterodimer. Interestingly we find that Mb expression which is driven by iron restriction and high sGC activation in iron deficiency anemia (IDA) leaches out more into the serum relative to non-anemic conditions. In line with our findings, analysis of tissues from IDA mice developed either by nutritional iron deprivation or by ablation of ferroportin (Fpn) gene or from IRIDA mice (caused due to TMPRRS6 KO), displayed a variable pattern of Mb expression in different tissues but it always correlated to the status of the sGC heterodimer or its subsequent activation, implying that Mb expression (either heme-free/apo-Mb or heme-containing/holo-Mb) is sGC dependent. We infer that Mb expression happening in anemic (IDA, Fe<5 ppm or IDA, Fpn) mice can be both due to iron restriction and an elevated sGC heterodimer that corroborates with greater sGC activation, and an associated effect of this is the Mb leaching out in the anemic blood.

Analyzing the Mb expression in the cardiac and skeletal muscles, we found a contrasting pattern of Mb expression (Fig. 4A, B) and was primarily due to opposing sGC heterodimer status in the two organs (Fig. 4D, E). While skeletal muscles from IDA (Fpn) mice showed a poor sGC heterodimer, cardiac IDA (Fpn) tissue displayed an elevated heterodimer that led to elevated Mb expression (Fig. 4B). The one difference in these two tissues is the absence of eNOS expression in the muscle cells (Fig. S1) while it is expressed in abundance in the vascular endothelial cells, which may cause a sustained sGC activation under IDA (Fpn) conditions in the cardiac tissue. This is similar to the lung tissues of IDA (Fe < 5ppm) mice which shows elevated sGC heterodimer relative to the non-anemic mice fed iron rich diet (Fe 350ppm) (Fig. 5B), and here as well there is abundant eNOS expression in the pulmonary artery endothelial cells or in the airway smooth muscle cells (33) of the lungs. Interestingly, the NO-sGC pathway seemed to be better activated in the IDA (Fpn) cardiac tissues or in the IDA (Fe < 5ppm) lung tissues relative to the WTs, as evidenced by an elevated sGC heterodimer (Fig. 4E and 5B) that displayed more than two-fold activation by NO and BAY-41 in IDA (Fpn) relative to WTs (Fig. 4F). However interesting these observations maybe, we currently cannot explain how sGC gets its heme to form an elevated sGC heterodimer under anemic conditions in an organ specific manner (eg. the heart and lungs). Whether HIF activation under IDA conditions (31, 40) can also increase sGC heterodimerization/activation is an intriguing possibility that remains to be determined.

While the current literature describes Rhabdomyolysis or Mb breakdown during injury, as a condition triggered by rapid skeletal muscle breakdown, causing heme-containing Mb to release in the blood and increasing toxicity before it is excreted out in the urine (41), our study suggests that an increase in heme-free Mb or apo-Mb in the muscle tissue can also be the genesis of Mb leaching into the blood serum. And this can be a molecular signature of Mb dysfunction in anemia or in any other diseased condition where this phenomenon is prevalent. Although studies suggest that iron homeostasis is vital for effective functioning of skeletal muscle, explicit studies linking muscle weakening to loss of Mb heme under anemic conditions were lacking (42). Moreover muscle weakening or muscle fatigue, a common symptom in anemia is primarily cited to be due to reduced oxygen delivery to muscles on account of low red blood cell count. This overlooks the additional fact that muscle weakening can also arise due to the inability of heme-free Mb to bind the bioavailable O_2_ thereby causing oxygen deprivation in the muscles. The genesis of this heme-free Mb primarily arises from a low or inactive sGC heterodimer in the myoblasts which can further block myoblast differentiation as realized from our studies (Fig. S1) (43), suggesting that Mb heme is also essential for muscle development. Our current study is probably the first study to date which sheds light on these heme-centric aspects and explains the anemic condition better by indicating that generation of heme-free Mb to be a critical event in anemic pathology and that the detected Mb leaching may be an associated outcome.

## Abbreviations

BAY 41: BAY 41-2272
BM: Bone Marrow
EPO: Erythropoietin
Hb: Hemoglobin
Mb: Myoglobin
Fpn: Ferroportin
IDA: Iron Deficiency Anemia
IRIDA: Iron Refractory Iron Deficiency Anemia
TMPRSS6: Transmembrane Serine Protease 6

## Data Availability Statement Included in the article

The data that support the findings of this study are available in the methods, results and/or supplementary material of this article.

## Acknowledgements

This work was supported by National Institute of Health Grants R01CA148828 (Y.S.), R01DK095201 (Y.S.), R01NS137230 (C.G.), R01GM148664 (D.J.S) and R01HL150049 (A.G.)

## Author Contributions

N. K. Das, M. P. Sumi, C. Ghosh, B. Tupta and A. Ghosh designed the experiments. N. K. Das, M. P. Sumi, B. Tupta, C. Ghosh and A. Ghosh performed all cell culture and biochemical studies. N. K. Das, M. P. Sumi, B.Tupta, C. Ghosh, D. J. Stuehr, Y. Shah and A. Ghosh analyzed all the data. A. Ghosh wrote the manuscript.

## REFERENCES

1. K. Chen, R. N. Pittman, A. S. Popel, Nitric oxide in the vasculature: where does it come from and where does it go? A quantitative perspective. Antioxid Redox Signal 10, 1185–1198 (2008).

2. R. F. Furchgott, J. V. Zawadzki, The obligatory role of endothelial cells in the relaxation of arterial smooth muscle by acetylcholine. Nature 288, 373–376 (1980).

3. R. M. Palmer, A. G. Ferrige, S. Moncada, Nitric oxide release accounts for the biological activity of endothelium-derived relaxing factor. Nature 327, 524–526 (1987).

4. R. C. Jin, J. Loscalzo, Vascular Nitric Oxide: Formation and Function. J Blood Med 2010, 147–162 (2010).

5. M. W. Radomski, R. M. Palmer, S. Moncada, Comparative pharmacology of endothelium-derived relaxing factor, nitric oxide and prostacyclin in platelets. Br J Pharmacol 92, 181–187 (1987).

6. J. Loscalzo, Nitric oxide insufficiency, platelet activation, and arterial thrombosis. Circ Res 88, 756–762 (2001).

7. D. I. Simon et al., Antiplatelet properties of protein S-nitrosothiols derived from nitric oxide and endothelium-derived relaxing factor. Arterioscler Thromb 13, 791–799 (1993).

8. C. M. Samama et al., Inhibition of platelet aggregation by inhaled nitric oxide in patients with acute respiratory distress syndrome. Anesthesiology 83, 56–65 (1995).

9. A. Ahluwalia et al., Antiinflammatory activity of soluble guanylate cyclase: cGMP-dependent downregulation of P-selectin expression and leukocyte recruitment. Proc Natl Acad Sci U S A 101, 1386–1391 (2004).

10. F. Murad, Shattuck Lecture. Nitric oxide and cyclic GMP in cell signaling and drug development. N Engl J Med 355, 2003–2011 (2006).

11. D. J. Stuehr, Mammalian nitric oxide synthases. Biochim Biophys Acta 1411, 217–230 (1999).

12. D. J. Stuehr, M. M. Haque, Nitric oxide synthase enzymology in the 20 years after the Nobel Prize. Br J Pharmacol 176, 177–188 (2019).

13. M. A. Marletta, Nitric oxide synthase: aspects concerning structure and catalysis. Cell 78, 927–930 (1994).

14. V. Kapil et al., The Noncanonical Pathway for In Vivo Nitric Oxide Generation: The Nitrate-Nitrite-Nitric Oxide Pathway. Pharmacol Rev 72, 692–766 (2020).

15. N. S. Bryan, K. Bian, F. Murad, Discovery of the nitric oxide signaling pathway and targets for drug development. Front Biosci (Landmark Ed) 14, 1–18 (2009).

16. E. R. Derbyshire, M. A. Marletta, Biochemistry of soluble guanylate cyclase. Handb Exp Pharmacol 10.1007/978-3-540-68964-5_2, 17–31 (2009).

17. E. R. Derbyshire, M. A. Marletta, Structure and regulation of soluble guanylate cyclase. Annu Rev Biochem 81, 533–559 (2012).

18. A. Ghosh, J. P. Stasch, A. Papapetropoulos, D. J. Stuehr, Nitric oxide and heat shock protein 90 activate soluble guanylate cyclase by driving rapid change in its subunit interactions and heme content. J Biol Chem 289, 15259–15271 (2014).

19. Ghosh et al., Low levels of nitric oxide promotes heme maturation into several hemeproteins and is also therapeutic. Redox Biol 56, 102478 (2022).

20. M. P. Sumi, B. Tupta, A. Ghosh, Nitric Oxide Trickle Drives Heme into Hemoglobin and Muscle Myoglobin. Cells 11 (2022).

21. L. Chen et al., Dynamic changes in murine erythropoiesis from birth to adulthood: implications for the study of murine models of anemia. Blood Adv 5, 16–25 (2021).

22. A. Kumar, E. Sharma, A. Marley, M. A. Samaan, M. J. Brookes, Iron deficiency anaemia: pathophysiology, assessment, practical management. BMJ Open Gastroenterol 9 (2022).

23. J. S. Souza, E. L. Brunetto, M. T. Nunes, Iron restriction increases myoglobin gene and protein expression in Soleus muscle of rats. An Acad Bras Cienc 88, 2277–2290 (2016).A

24. A. Ghosh, Y. Dai, P. Biswas, D. J. Stuehr, Myoglobin maturation is driven by the hsp90 chaperone machinery and by soluble guanylyl cyclase. FASEB J 33, 9885–9896 (2019).

25. M. Dziegala et al., Iron deficiency as energetic insult to skeletal muscle in chronic diseases. J Cachexia Sarcopenia Muscle 9, 802–815 (2018).

26. K. E. Finberg et al., Mutations in TMPRSS6 cause iron-refractory iron deficiency anemia (IRIDA). Nat Genet 40, 569–571 (2008).

27. Y. Yu et al., Activation of Intestinal HIF2alpha Ameliorates Iron-Refractory Anemia. Adv Sci (Weinh) 11, e2307022 (2024).

28. Y. Zhao et al., Inhibition of soluble guanylate cyclase by ODQ. Biochemistry 39, 10848–10854 (2000).

29. Tupta et al., GAPDH is involved in the heme-maturation of myoglobin and hemoglobin. FASEB J 36, e22099 (2022).

30. N. K. Das et al., Microbial Metabolite Signaling Is Required for Systemic Iron Homeostasis. Cell Metab 31, 115–130 e116 (2020).

31. A. J. Schwartz, K. Converso-Baran, D. E. Michele, Y. M. Shah, A genetic mouse model of severe iron deficiency anemia reveals tissue-specific transcriptional stress responses and cardiac remodeling. J Biol Chem 294, 14991–15002 (2019).

32. J. Schwartz et al., Hepatic hepcidin/intestinal HIF-2alpha axis maintains iron absorption during iron deficiency and overload. J Clin Invest 129, 336–348 (2019).

33. M. P. Sumi et al., Expression of soluble guanylate cyclase (sGC) and its ability to form a functional heterodimer are crucial for reviving the NO-sGC signaling in PAH. Free Radic Biol Med 225, 846–855 (2024).

34. A. G. Pinder, S. C. Rogers, A. Khalatbari, T. E. Ingram, P. E. James, The measurement of nitric oxide and its metabolites in biological samples by ozone-based chemiluminescence. Methods Mol Biol 476, 11–28 (2008).

35. Hausladen et al., Assessment of nitric oxide signals by triiodide chemiluminescence. Proc Natl Acad Sci U S A 104, 2157–2162 (2007).

36. Ghosh, G. Garee, E. A. Sweeny, Y. Nakamura, D. J. Stuehr, Hsp90 chaperones hemoglobin maturation in erythroid and nonerythroid cells. Proc Natl Acad Sci U S A 115, E1117–E1126 (2018).

37. R. Liu et al., Heme oxygenase 1 in erythropoiesis: an important regulator beyond catalyzing heme catabolism. Ann Hematol 102, 1323–1332 (2023).

38. S. T. Fraser, R. G. Midwinter, B. S. Berger, R. Stocker, Heme Oxygenase-1: A Critical Link between Iron Metabolism, Erythropoiesis, and Development. Adv Hematol 2011, 473709 (2011).

39. S. T. Fraser et al., Heme oxygenase-1 deficiency alters erythroblastic island formation, steady-state erythropoiesis and red blood cell lifespan in mice. Haematologica 100, 601–610 (2015).

40. Y. M. Shah, L. Xie, Hypoxia-inducible factors link iron homeostasis and erythropoiesis. Gastroenterology 146, 630–642 (2014).

41. J. F. Hebert, K. G. Burfeind, D. Malinoski, M. P. Hutchens, Molecular Mechanisms of Rhabdomyolysis-Induced Kidney Injury: From Bench to Bedside. Kidney Int Rep 8, 17–29 (2023).

42. M. Stugiewicz et al., The influence of iron deficiency on the functioning of skeletal muscles: experimental evidence and clinical implications. Eur J Heart Fail 18, 762–773 (2016).

43. J. S. J. Vinke et al., Iron deficiency is related to lower muscle mass in community-dwelling individuals and impairs myoblast proliferation. J Cachexia Sarcopenia Muscle 14, 1865–1879 (2023).

