## Supplementary Material for "Myoglobin leaching into the serum of IDA mice is driven by the high activation of sGC under anemic conditions which induces myoglobin expression"

**Supplementary Material: Tables S1 and Figure S1 with Figure legends**

**Table S1. Antibodies used and their sources.**

**Figure S1. Impact of sGC inhibition on myoblast differentiation and Mb heme-insertion.** Mouse C2C12 myoblasts were triggered for differentiation (0-96 h) with culture media containing 2% horse serum, while parallel cultures were treated with sGC inhibitor ODQ between 24-96h to study the impact on differention and Mb heme-maturation. (A) Expression of NOSs, differentiation marker & Mb heme-stain during C2C12 differentiation and impact of sGC inhibition (by ODQ) on differentiation. (B) IPs showing status of sGC$\alpha 1\beta 1$ heterodimer during differentiation, -/+ ODQ and correlation of Mb heme densitometries as shown in panel A plotted against sGC heterodimer status obtained from the IPs depicted in panel B. (C) Nitrite levels from NO generated by NOSs during myoblast differentiation, -/+ ODQ. Data are mean -/+ SD of n=3 experiments.

**Tables S1. Antibodies used and their sources.**

| Serial No. | Antibody | Species-Clonality | Source (Catalog No.) |
|---|---|---|---|
| 1. | β-Actin | Mouse monoclonal | SIGMA (A5441) |
| 2. | eNOS | Rabbit polyclonal | Cell Signaling Tech. (9572) |
| 3. | iNOS | Mouse monoclonal | Invitrogen (MA5-17139) |
| 4. | nNOS | Rabbit polyclonal | Invitrogen (PA3-032A) |
| 5. | EPO | Mouse monoclonal | Santa Cruz (sc-5290) |
| 6. | HO1 | Rabbit polyclonal | Cell Signaling Tech. (70081) |
| 7. | Myogenin | Rabbit monoclonal | Abcam (ab124800) |
| 8. | Myoglobin | Mouse monoclonal | Santa Cruz (sc-393020) |
| 9. | sGCβ1 | Rabbit polyclonal | SIGMA, ER-19 (G4405) |
| 10. | sGCβ1 | Rabbit polyclonal | Cayman Chem. (160897) |
| 11. | sGCα1 | Rabbit polyclonal | Abcam (ab101368) |

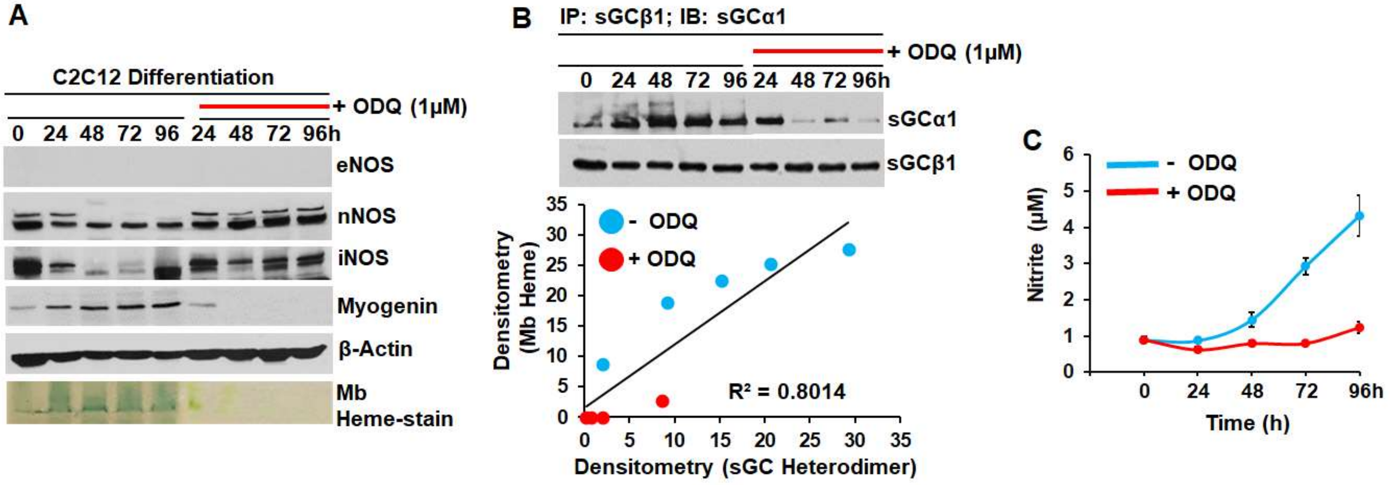
A
C2C12 Differentiation
+ ODQ (1μM)
0 24 48 72 96 24 48 72 96h
eNOS
nNOS
iNOS
Myogenin
β-Actin
Mb
Heme-stain
B
IP: sGCβ1; IB: sGCα1
+ ODQ (1μM)
0 24 48 72 96 24 48 72 96h
sGCα1
sGCβ1
- ODQ
+ ODQ
R² = 0.8014
Densitometry (Mb Heme)
Densitometry (sGC Heterodimer)
C
- ODQ
+ ODQ
Nitrite (μM)
Time (h)
0 24 48 72 96h

**Figure S1.**
